# Mitochondria induce anisotropy and delays in action potential conduction

**DOI:** 10.1101/2023.03.28.534468

**Authors:** Ann Castelfranco, Pepe Alcami

## Abstract

The internal resistance of axons to ionic current flow affects the speed of action potential propagation. As biological cables, axons contain mitochondria which are necessary to support axonal function with energy supply. Although we would expect mitochondria to increase the internal resistance to current flow, their impact on the conduction velocity of action potentials has remained elusive. To investigate the impact of mitochondria on action potential propagation in the small non-myelinated fibers found in the vertebrate brain, we combined computational modeling and electron microscopy from the axons found in the premotor pathway that controls the production of birdsong with submillisecond precision. Mitochondria occupancy of axonal cross-sections ranged from 5 to 73% (average: 29%) in the ∼ 0.2-0.7 μm diameter non-myelinated axons connecting song premotor nuclei HVC and RA in canaries. Interestingly, this occupancy depends on axonal diameter: axonal cross-section occupancy by mitochondria was larger in small axons, with an average occupancy of ∼46% for axons with diameters smaller than 300 nm and ∼21% for larger diameters. Computational modeling showed that when the propagating action potential meets a mitochondrion, the conduction velocity decreases and the action potential is delayed by tenths of microseconds to microseconds. This effect is stronger in small axons given their larger cross section mitochondrial occupancy and cumulates delays of tenths of milliseconds along the whole pathway linking HVC and RA. Finally, we modeled the impact of varying densities of mitochondria on action potential propagation along the songbird premotor pathway. In summary, our model shows that axonal mitochondria induce the anisotropic propagation of action potentials, and that this effect cumulates a typical delay in the order of tenths of milliseconds over distances of mms. By partially occupying axoplasm, mitochondria constitute a biological design constraint that delays information processing in the small-diameter unmyelinated axons found in the vertebrate brain.

## Materials and methods

### Animals

Three adult male canaries (Serinus canaria) housed in outdoor aviaries at the Max Planck Institute for Ornithology (Seewiesen) were euthanized by an overdose of isoflurane (two for electron microscopy and one for light microscopy). Housing, welfare of the animals and experimental procedures complied with the requirements of the European Directives for the protection of animals used for scientific purposes 2010/63/EU of the European parliament, the German ‘Verordnung zum Schutz von zu Versuchszwecken oder zu anderen wissenschaftlichen Zwecken verwendeten Tiere’ and the German Animal Protection Act.

### Fixation

#### For electron microscopy

After death had been confirmed, animals underwent intracardiac perfusion for 2 minutes with a PBS solution containing sodium nitroprusside (VWR chemicals, 10µg/ml) followed by 20 minutes with a ‘Karlsson-Schultz’ perfusion solution containing 4% formaldehyde (Carl Roth Art.Nr. 0335), 2.5% glutaraldehyde (Electron Microscopy Sciences, cat.# E16220), 0.5% NaCl in phosphate buffer adjusted to pH 7.4 (Möbius et al. 2010). Perfusion speed was 1ml/min. Brains were postfixed for 24h.

#### For light microscopy

After death had been confirmed, the canary underwent intracardiac perfusion for 2 minutes with a PBS solution containing sodium nitroprusside (VWR chemicals, 10µg/ml) followed by 20 minutes with a perfusion solution containing 4% formaldehyde (Carl Roth Art.Nr. 0335) in PBS. Perfusion speed was 1ml/min. The brain was postfixed for 24h.

### Electron microscopy

#### Vibratome sections

Sagittal sections 100 µm to 300 µm thick were sliced with a vibratome (Leica VT1200S). Sections approximately 1 µm by 1 µm isolated from the region containing bundles that exit the nucleus HVC in the direction of RA were further processed for electron microscopy.

#### Osmification

Osmification was performed with 1% Osmium Tetroxide (2% Osmium Tetroxide, Electron Microscopy Sciences, cat.#19152) in 0.1M Sodium Cacodylate pH 7.4 (Sodium Cacodylate buffer 0.2 M, Electron Microscopy Sciences, cat.#11653), for 40 min.

#### Dehydration

Osmification was followed by 3 rounds of washing in distilled water and dehydration in successive steps, each of 10 minutes, in 30%, 50%, 70%, 100% Ethanol for 10 minutes. Samples were embedded in Spurr’s low viscosity embedding medium (Electron Microscopy Sciences, cat.#14300) according to the manual (https://www.emsdiasum.com/docs/technical/datasheet/14300) for 48h at 60 °C.

#### Semi-thin sections

Slices were cut at 0.5 µm thickness with the ultramicrotome EM UC6 (Leica) and stained with epoxy tissue stain (Electron Microscopy Sciences, cat.#14950).

#### Ultra-thin sections

Slices were cut at 60 nm thickness with ultramicrotome EM UC6 (Leica). *Contrast counterstain*. Sections were stained with ‘ultrostainer’ (Leica) with 0.5% uranyl acetate (Uranyl acetate solution 1%, Electron Microscopy Sciences, cat.#22400-1) and 3% lead citrate (Ultrostain 2, Leica).

#### Image acquisition

Images were acquired with a JEOL (JEM-1230) transmission electron microscope and a Gatan Orius SC1000 digital Camera with the software Gatan DigitalMicrograph™.

#### Image analysis

Quantifications were performed on magnifications of 40.000 to 80.000 from axon bundles between HVC and RA. Images were analyzed with ImageJ software (https://imagej.net/software/fiji/) and Igor pro software (https://www.wavemetrics.com/). We fitted the axons and mitochondria to ellipses with ImageJ built-in plugin. Occupancy of axonal cross-sections by mitochondria was measured as the ratio of the area estimated from a disk whose diameter was set to the minor axis of an ellipse fitted to the mitochondrion to that analogously estimated for the corresponding axon. In three axon cross sections we found two mitochondria. These were excluded from the mitochondria to axon ratio calculations based on the ellipse fits.

The total volumetric occupancy was estimated from the measurement of the volume occupied by non-myelinated axons inside a bundle in longitudinal sections of bundles. Large fractions of extracellular space found in the bundle were deducted from this volume.

### Confocal microscopy

#### Vibratome sections

Brain sagittal sections 60 µm thick were sliced with a vibratome (Leica VT1200S).

#### Immunostaining

A sagittal slice containing nuclei HVC and RA was incubated in a blocking solution (BS) containing bovine 1% serum albumin (weight/volume), 0.1% saponin and 1% tritonX-100 in PBS at room temperature for 1h. The slice was then incubated with the primary antibody against the Neurofilament heavy chain, NFH (Abcam ab4680, chicken polyclonal IgY, dilution 1:400 in BS) at 4°C for 48h, washed four times in BS at room temperature, and incubated in AMCA anti-chicken secondary antibody (Dianova, 703-156-155, dilution 1:200 in BS) at 4°C for 24h, washed once in BS and three times in PBS and mounted in Vectashield mounting medium.

#### Image acquisition

Acquisitions were performed at the Center for Advanced Light Microscopy (LMU) with a Ti-E Nikon spinning disk microscope equipped with a CFI Apochromat LWD Lambda S 40XC WI objective and an Andor iXon Ultra 888 EMCCD camera. The fluorochrome was visualized with an excitation wavelength of 405 nm (emission filter 420-460nm). Images were acquired with a pixel size of 326 nm and averaging four planes.

### Computational modeling

Models for the unmyelinated axons of HVC projection neurons that run in bundles to the premotor nucleus RA (HVC_RA_ cells) in canaries were simulated using the NEURON simulation environment (version 7.4) (Carnevale and Hines 2006; Hines and Carnevale 1997). Based on the morphology of these axons, the simple ball and stick model neuron consisted of a spherical soma 6 µm in diameter with a single cylindrical axon less than 1 µm in diameter. Simulated axon diameters ranged from 0.1 µm to 0.7 µm, in agreement with experimental measurements reported here. The soma contained only a passive leak conductance while the axons contained fast sodium and delayed rectifier potassium conductances in addition to the leak conductance. The descriptions of the ion channel kinetics were taken from the model for mammalian neocortical pyramidal axons of (Cohen et al. 2020) available from the ModelDB database (https://senselab.med.yale.edu/ModelDB/showmodel?model=260967). The fast sodium channel has the 8-state kinetic gating scheme of (Schmidt-Hieber and Bischofberger 2010). The potassium channel kinetics were described using the Hodgkin-Huxley formalism (Hodgkin and Huxley 1952) for a non-inactivating potassium channel with parameters based on a Kv1.1 subunit (Akemann and Knöpfel 2006). The ion channel densities were set to 1000 pS/cm^2^ for the sodium channel and 3000 pS/cm^2^ for the potassium channel and were uniform along the axon. These channel densities were chosen to fit the amplitude of an action potential recorded from a canary HVC_RA_ cell at 20 ºC (unpublished observation) and the range of conduction delays measured from HVC to the RA along axons putatively identified as unmyelinated (Hahnloser, Kozhevnikov, and Fee 2006; Egger et al. 2020). The simulations were run at 40 ºC according to physiological canary body temperatures. The cytoplasmic resistivity of the axon without mitochondria was set to 100 Ωcm, while that of the sections containing a mitochondrion was varied to model the effect of the mitochondrion (see Results). For all sections, the specific membrane capacitance was set to 1µF/cm^2^, and the reversal potential of the leak current to -70 mV.

The axon was made up of two types of sections: those containing a mitochondrion and those with just axoplasm. Generally, the mitochondrion-containing compartments were 1 µm in length and were distributed uniformly along the axon. The effect of varying the length and distribution of the mitochondrial compartments on the average computed action potential conduction velocity was usually small as long as the “total amount” of mitochondria in the axon remained fixed. In particular, when the 1 µm mitochondrion compartments were distributed randomly instead of uniformly, the relative difference in the conduction velocity was less than 0.2%. However, in the extreme case when very large mitochondria were collected in one long compartment, the relative change in conduction velocity was ∼11%. Conduction velocity was computed from the difference between the times when the action potential upswing crossed -5 mV at two positions near the middle of the axon separated by a known distance, typically 200 µm. The time when the membrane potential crossed -5 mV was interpolated from the times of adjacent points on the voltage trajectory spanning -5 mV. To compute the local effect on conduction velocity of a single mitochondrion the simulation was run with all axon compartments containing mitochondria.

The simulations used NEURON’s default backward Euler integration with a time step size of 2.5 µs. The soma was subdivided into 2 µm compartments; the spatial grid for the axon was finer, typically 0.33 µm and 0.82 µm compartments for axon sections with and without mitochondria, respectively. A 3-fold increase in the fineness of the spatial grid resulted in a relative change in the conduction velocity of less than 0.01%. Action potentials were elicited by a 0.5 ms duration 0.5 nA current pulse to the soma following a 10 ms delay to allow any membrane voltage transients to settle back to the resting membrane potential.

## Introduction

The timely propagation of action potentials requires the fine tuning of axonal biophysical properties to ensure the correct functioning of neural circuits (Rushton 1951; Deutsch 1969; Seidl, Rubel, and Harris 2010; Castelfranco and Hartline 2015; Alcami and El Hady 2019). A fundamental functional property of action potential propagation, conduction velocity, depends on the morphological and physiological properties of axons and, when myelinated, of their myelinating cells. Axonal morphology constrains the passive properties of axons, a phenomenon that is today well understood (Manor, Koch, and Segev 1991; Ofer, Shefi, and Yaari 2020). However, a property of biological cables, and its impact on conduction speed, has remained elusive: the presence of intracellular organelles. One such prevalent organelle found in axons are mitochondria.

In comparison to non-biological cables, biological cables such as axons require the supply of energy all along the cable to maintain their function (Perge et al. 2009; Harris and Attwell 2012; Sterling and Laughlin 2015). Mitochondria provide axons with energy, required among others to mantain resting potentials and propagate action potentials, housekeeping and synaptic functions (Harris and Attwell 2012). By occupying intracellular volume, we would expect that mitochondria reduce cytosolic cross-sectional area and thereby increase the axial resistance to current flow and the conduction velocity of propagating action potentials. However, their impact on action potential propagation remains elusive, and axons have so far been modeled as cables without organelles (Meunier and Segev 2001).

In this article, we investigate the impact of mitochondria on action potential conduction velocity in the characteristic small unmyelinated axons found in vertebrate brains, with diameters in the range of hundreds of nanometers (Perge et al. 2009; Braitenberg 1991; Wang et al. 2008), i.e. (Braitenberg 1991) measured an average axonal diameter below 0.3 μm in the mammalian neocortex, and (Aboitiz et al. 1992) found corpus callosum unmyelinated axon diameters to be in the range of 0.1 to 1 μm. We focus on the premotor pathway involved in the fast control of vocal muscles that produce birdsong, that is, the axons belonging to the principal cells of the song nucleus HVC (formerly known as ‘high vocal center’, used here as proper name) that project to the RA (robust nucleus of the arcopallium), the HVC_RA_ cells. HVC_RA_ cells fire action potentials with submillisecond precision and in high-frequency bursts during singing (Hahnloser, Kozhevnikov, and Fee 2002). Thus, this pathway is a model of choice to investigate delays in action potential propagation that could be induced by mitochondria.

## Results

### Mitochondrial occupancy of unmyelinated axons depends on axonal diameter

We examined with transmission electron microscopy the pathway connecting the two premotor regions, the ‘song nuclei’ involved in birdsong production, HVC and RA (Fig. 1A) in the canary (*Serinus canaria*). This motor pathway is formed by a mixture of myelinated and unmyelinated axons, spatially clustered together in axon bundles (Fig. 1B).

**Figure 1.**
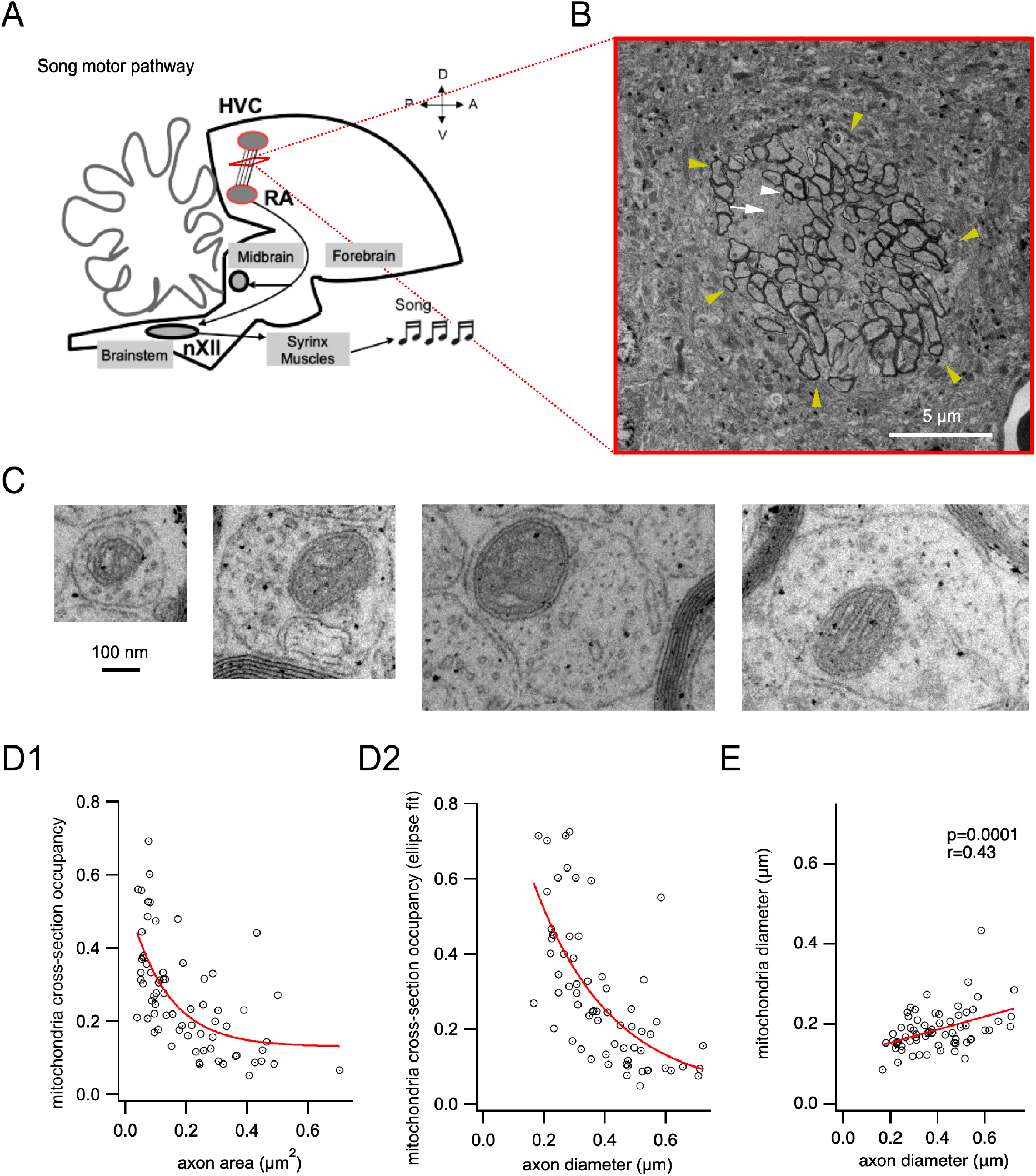
Mitochondria occupancy of unmyelinated axonal cross-sections between nuclei HVC and RA decreases with axon diameter. **A**. Schematic representation of the axons running between HVC and RA in the song motor pathway. The song motor pathway controlling birdsong production originates in HVC within the nidopallium, continues in downstream nucleus RA in the arcopallium, whose neurons project onto nucleus nXII in the brainstem, which in turn projects to the vocal muscles that form the songbird vocal organ, the syrinx. Modified from (Alcami et al. 2021). **B**. Example of axon bundle (yellow arrows) formed by axons from the pathway between HVC and RA. Axons comprise myelinated axons (white arrowhead) and unmyelinated axons (white arrow). **C**. Representative examples of axonal cross-sections of different sizes containing mitochondria, imaged with transmission electron microscopy. **D1**. Occupancy of axonal cross-sections by mitochondria, measured as the ratio of mitochondrial to axonal areas, plotted as a function of measured axon area. An exponential fit of the data is shown in red. **D2**. Occupancy of axonal cross-sections by mitochondria, measured as the ratio of the area estimated from a disk whose diameter was set to the minor axis of an ellipse fitted to the mitochondrion to that analogously estimated for the corresponding axon, as a function of axon diameter. An exponential fit of the data is shown in red. **E**. Mitochondrion diameter plotted as a function of axon diameter. A linear fit of the data is plotted in red. The p and r values from a linear correlation test are indicated.

We measured the fraction of the non-myelinated axonal cross-sectional area occupied by a mitochondrion from mitochondrion-containing axonal radial sections, which we termed the “cross-sectional mitochondrial occupancy” of the axon. Representative examples of axons of different sizes are shown in Fig. 1C. The cross-sectional mitochondrial occupancy ranged from 5.2% to 69.3% (Figure 1D1), with an average of 26.4 ± 1.8% (n = 69 cross-sections of axons containing mitochondria from two canaries). Alternatively, we fitted ellipses to mitochondria and axons and calculated areas based on the minor axis of the ellipse, assuming the axons were cylindrical, correcting for planes not perfectly orthogonal to axon bundles (Harris and Attwell 2012). This quantification gave similar results, with fractions ranging from 4.8% to 72.5% and averaging 29.1 ± 2.3% (Fig. 1D2).

Non-myelinated axon diameters spanned the range from 166 nm to 724 nm (n = 66), averaging 394 ± 17 nm. In order to examine whether cross-sectional mitochondrial occupancy varied depending on axon diameter, we plotted it as a function of axon diameter (Fig.1D2). Interestingly, cross-sectional mitochondrial occupancy, both calculated from area measurements on electron micrographs and based on ellipse fits, showed a negative correlation with axon area and diameter (Fig. 1D1-D2, linear correlation test, p = 9*10 ^-8^, r = - 0.59; and p = 3*10^−9^, r = -0.65 respectively). That is, a larger fraction of the axon cross-section was covered in small axons relative to larger axons. Whereas on average mitochondrial occupancy accounted for 46.2% of the axon cross-section (by the fitted ellipse method) for axons up to 300 nm in diameter, this fraction decreased to 21.2% for axons larger than 300 nm in diameter.

These results suggested little or no scaling of mitochondria with axon size. Indeed, we confirmed that the distributions of mitochondria and axon diameters showed a small correlation (Fig.1E), which could be fitted by a linear function with slope 0.163 (p = 0.0001, r = 0.43).

### Model for mitochondrion-containing axonal sections

To model the impact of a mitochondrion on conduction velocity, we made the simplifying assumption that the mitochondrion acts like a region of high resistance to axial current flow. Consider a section of an axon that contains a cylindrical mitochondrion. Since the mitochondrion doesn’t completely fill the cross-section of the axon, there are two current paths through the core of the cylindrical axon section: one passes through the cytoplasm avoiding the mitochondrion and the other passes through the mitochondrion (Fig. 2C1). This can be described by a circuit with two impedances in parallel, which can be approximated (Padmaraj et al. 2014) by two resistors in parallel, where *r*_*ax*_ (Ω) is the resistance of the path through the cytoplasm and *r*_*mit*_ (Ω) is the resistance of the path through the mitochondrion (Figure 2C2). These resistances can be replaced by an equivalent resistance *r*_*eq*_ given by:

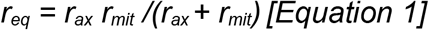

Rewriting the equivalent resistance, *r*_*eq*_, in terms of the intracellular resistivity, *R*_*eq*_ (Ω cm), gives:

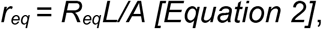

where *A* is the cross-sectional area of the cylindrical axon and *L* is the length of the section containing the mitochondrion, which we define to be the length of the mitochondrion.

**Figure 2.**
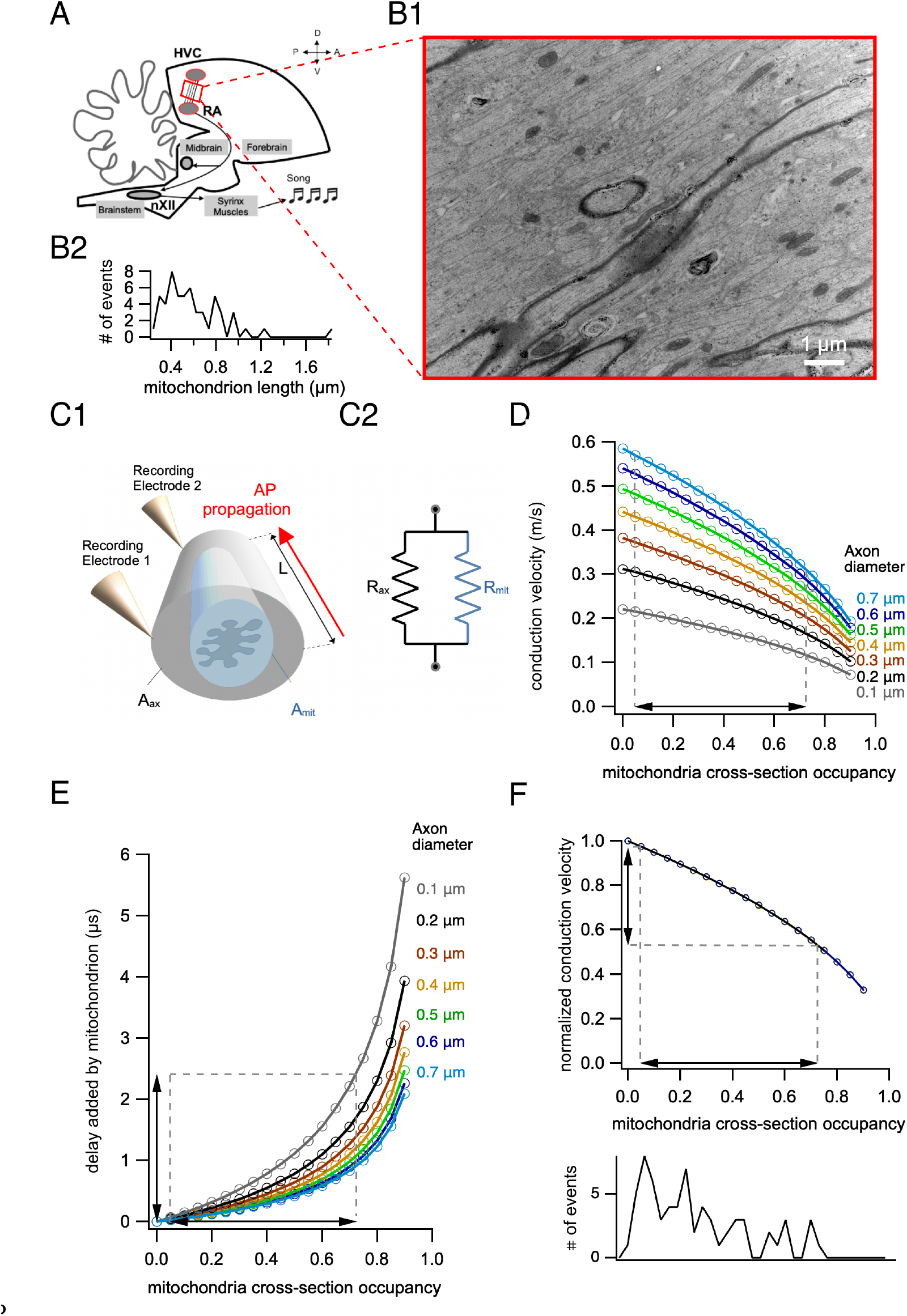
Impact of a single mitochondrion on action potential conduction velocity. **A**. Schematic of the pathway between HVC and RA. Modified from (Alcami et al. 2021). **B**. Longitudinal section of an axon bundle. Mitochondria length estimated from the major axis of an ellipse fit. **C1**. Schematic of the modeled section of an axon. Action potential propagation was modeled along a cylinder of 0.6 μm in length (L) and conduction speed measured between the two extremes of the cylinder. **C2**. Equivalent circuit of the axial component used in our model. **D**. Action potential conduction velocity decreases for cylinders containing mitochondria relative to an axon free of mitochondria, as a function of cross-sectional mitochondrial occupancy of the axon for different axon diameters. The range of cross-section occupancies measured in our data is indicated with a dashed line. **E**. Action potential latency increases for cylinders containing mitochondria relative to an axon free of mitochondria, as a function of cross-sectional mitochondrial occupancy of the axon. The range of cross-section occupancies measured in our data is indicated with a dashed line. **F**. Top, the same functions normalized to a mitochondria-free axon overlapped for all diameters. Bottom, histogram of measured cross-sectional occupancies. The correspondiong range is marked with dashed lines on the top graph.

The cross-sectional area, *A*, can be partitioned into the area taken up by the mitochondrion, *A*_*mit*_ and the area free from the mitochondrion, *A*_*ax*_.

Let *p* be the proportion of *A* that is taken up by the mitochondrion, then:

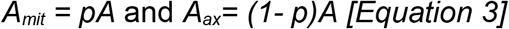

Combining equations 2 and 3 gives:

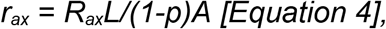

where *R*_*ax*_ is the cytoplasmic resistivity of the axon and similarly,

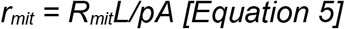

Hence, combining equations 1, 4 and 5, the equivalent resistivity of the section is:

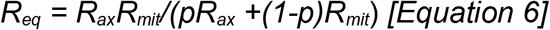

Thus, given estimates for the intracellular resistivity of the axon and the resistivity of a mitochondrion, we can estimate the combined resistivity in terms of the cross-sectional mitochondrial occupancy.

The resistivity of a mitochondrion is not a particularly well-defined quantity, but we can make a rough estimate for a model cylindrical mitochondrion 0.2 µm in diameter (Fig. 1E) and 1 µm in length. If we assume that the resistance across a unit area of mitochondrial outer membrane is similar to that of the axolemma, on the order of 10^4^ Ωcm^2^, then for a cylindrical mitochondrion 0.2 µm in diameter the membrane resistance per unit length is ∼1.6×10^8^ Ωcm (Jack, Noble, and Tsien 1975). So the resistance of a 1 µm long mitochondrion is ∼1.6×10 ^12^Ω. Thus the resistivity of the model mitochondrion is approximately 5 MΩcm. This is a crude estimate that ignores much of the structure of the mitochondrion. However, (Jonas, Buchanan, and Kaczmarek 1999) using a patch clamp technique to record from mitochondria in the presynaptic terminal of the squid found only a small conductance of ∼28pS in the quiescent terminal. This corresponds to a membrane resistance of ∼4×10^10^ Ω for the patch. In order to use this resistance to estimate the resistivity, we need the dimensions of the resistor. The diameter of the tip of the patch electrode was ∼0.2 µm, but whether the recording was from the surface of the mitochondrion or extended to inner membranes was unclear. Assuming that this thickness was at most 1 µm, although likely to be much less, this gives an estimated resistivity of at least ∼10^5^ Ωcm. Hence, for the the model, we used the conservative estimate of 10^4^ Ωcm for the resistivity of a mitochondrion and 100 Ωcm for the intracellular resistivity of the axon.

### Mitochondria induce a local decrease in action potential conduction velocity

We used the computational model to investigate the impact of the cross-sectional mitochondrial occupancy on action potential propagation in unmyelinated axons. Since conduction velocity (Hodgkin and Huxley 1952) and cross-sectional mitochondrial occupancy (Fig.1) depend on axon diameter, we simulated action potential propagation for different axon diameters. We first explored the local effect of a single mitochondrion on conduction velocity.

In order to measure mitochondrial length on the longitudinal axis, we prepared longitudinal sections from axon bundles (Fig. 2. A, B1). The longitudinal length of a mitochondrion ranged from 242 to 1882 nm measured as the major axis from an ellipse (average: 622 ± 38 nm, n = 55, Fig. 2B2). Thus we report the local impact of a mitochondrion on action potential conduction for a typical mitochondrion length of 0.6 µm (Fig.3A,B).

**Figure 3.**
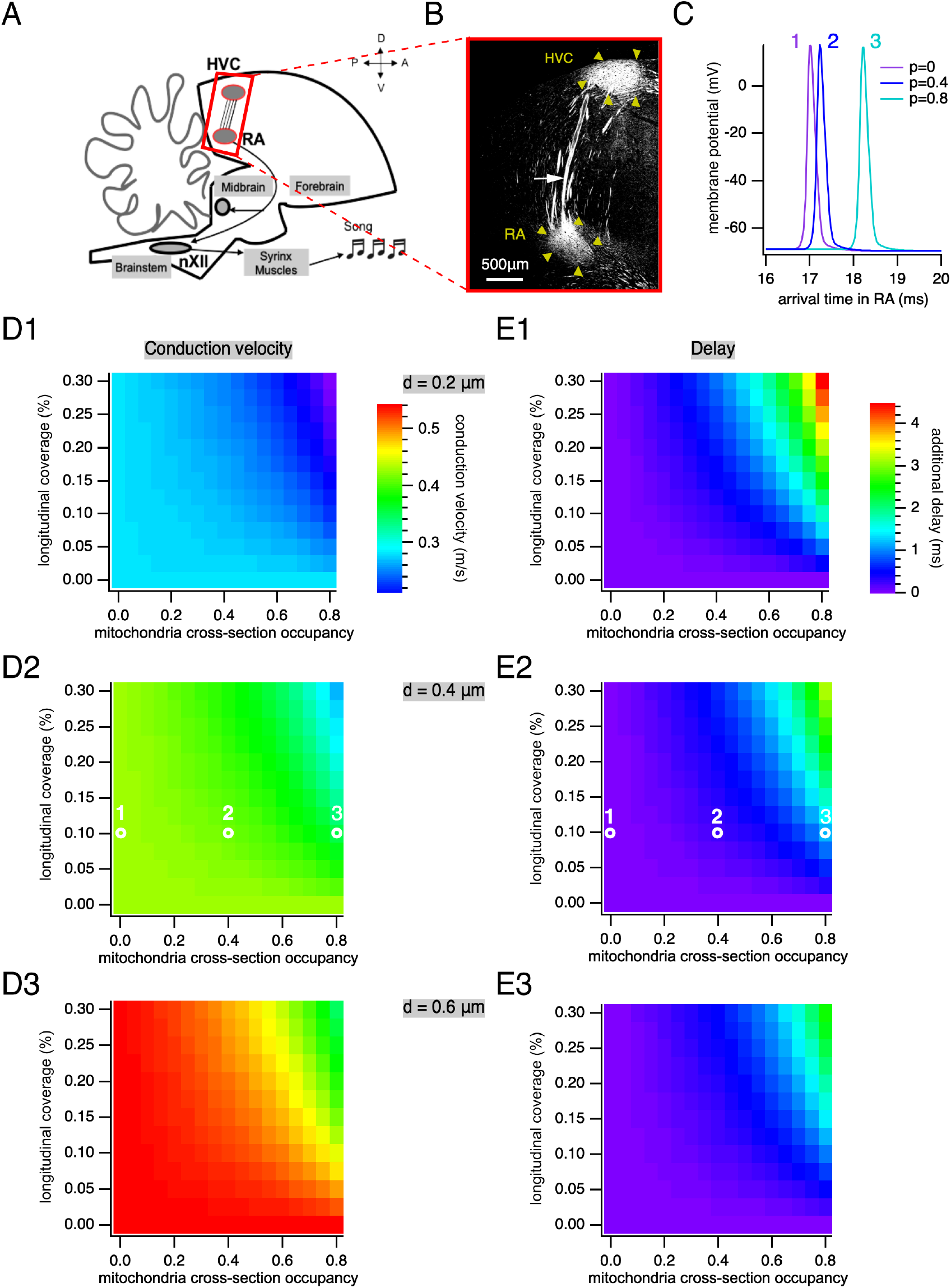
Action potential propagation between HVC and RA and influence of mitochondria density. **A**. Schematic of the pathway along which we modeled the propagation of action potentials, between HVC and RA. Modified from (Alcami et al. 2021). **B**. Immunostaining against the neurofilament heavy chain, which makes it possible to visualize the pathway formed by axons running in bundles between HVC and RA (both indicated by yellow arrowheads). The white arrow indicates an axon bundle. **C**. Example showing action potentials after stimulus onset with varying mitochondrial occupancy in 0.4 μm diameter axons with 10% longitudinal coverage at the RA (p = 0 in purple, p = 0.4 in dark blue and p = 0.8 in turquoise). **D1**. Action potential conduction velocity as a function of longitudinal coverage and mitochondria cross-section occupancy for axons of a diameter of 0.2 μm. **D2**. Same as D1 for 0.4 μm. White circles indicate the parameters of action potentials shown in C. **D3**. Same as D1 for 0.6 μm. **E1**. Additional delay of action potential arrival in RA a function of longitudinal coverage and mitochondria cross-section occupancy for axons of a diameter of 0.2 μm. **E2**. Same as E1 for 0.4 μm. White circles indicate the parameters of action potentials shown in C. **E3**. Same as E1 for 0.6 μm.

We compared the conduction velocity of an action potential propagating along the membrane of a 0.6 μm long cylindrical axon containing a mitochondrion (Fig. 2C1, D) with that of the same model axon devoid of mitochondria. Local conduction velocity decreased with increasing mitochondrial occupancy and decreasing axon diameter. For different typical cross-sectional occupancies and axon diameters, conduction velocity decreased by ∼ 0.11 m/s (36%) for small 0.2 μm-diameter axons with 60% cross-sectional occupancy, by ∼ 0.06 m/s (13%) for medium 0.4 μm-diameter axons with 25% cross-sectional occupancy and by ∼0.04 m/s (8%) for larger 0.6 μm-diameter axons with 15% cross-sectional occupancy. Thus, smaller axons are more susceptible to a decrease of conduction velocity induced by mitochondria.

We then computed the additional propagation delay through the mitochondrion-containing section for different axon diameters (Fig. 2E). Conduction delay increased with increasing mitochondrial occupancy and decreasing axon diameter. For typical cross-sectional mitochondrial occupancy in the measured biological range (Fig. 1), the additional delay caused by a single mitochondrion was ∼1.1 μs for small 0.2 μm-diameter axons with 60% cross-sectional occupancy, ∼ 0.2 μs for medium 0.4 μm-diameter axons with 25% cross-sectional occupancy and ∼ 0.1 μs for larger 0.6 μm-diameter axons with 15% cross-sectional occupancy.

Although conduction velocity for a given cross-sectional mitochondrial occupancy was larger for larger diameter axons (Fig. 2D), the relative decrement in velocity due to cross-sectional mitochondrial occupancy was the same for different axon diameters (Fig. 2F).

That is, when conduction velocity was normalized to that of the mitochondrion-free axon for each diameter, the relative decrease of conduction velocity was similar for all the diameters. This observation is consistent with the prediction that the conduction velocity of an unmyelinated axon is proportional to its *(diameter)*^*½*^, so for two axons 1 and 2 with the same underlying biophysical properties:

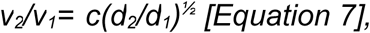

where *v*_*1*_, *d*_*1*_ and *v*_*2*_, *d*_*2*_ are the conduction velocity and diameter of axon 1 and axon 2, respectively and *c* is a constant (Hodgkin and Huxley 1952). Thus, for the same coverage for two axons with different velocities, their velocity ratio would be scalable by a constant *c(d*_*2*_*/d*_*1*_)^*½*^. In other words, the impact of mitochondrial coverage relative to the mitochondria-free axon normalizes to the conduction speed for each axon size, and is therefore comparable in relative magnitude.

### Longitudinal coverage of axons by mitochondria

The overall impact of mitochondria on the conduction velocity along the pathway will depend on the fraction of the axon length that contains mitochondria. We estimated the average longitudinal coverage of the axon by mitochondria used in our simulations from electron micrographs with the following equation:

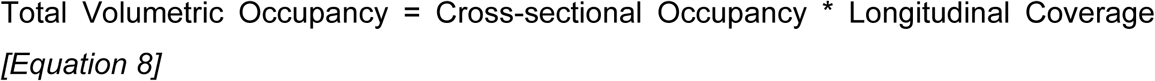

where *total volumetric occupancy* is the total fraction of the volume of non-myelinated axons in the micrograph occupied by mitochondria, *longitudinal coverage* is the fraction of axon length covered by mitochondria in the longitudinal section and *cross-sectional occupancy* is the fraction of axonal cross-section area covered by mitochondria as measured in Fig.1.

Hence, the total volumetric occupancy is proportional to the cross-sectional mitochondrial occupancy and to the average longitudinal mitochondrial coverage.

We measured the fraction of the non-myelinated fibers occupied by mitochondria in an electron micrograph from a longitudinal bundle (185.50 μm^2^ area *0.06 thickness), the resulting total volumetric occupancy equalled 0.0377, that is, ∼ 3.8%. Hence, the total occupancy in HVC_RA_ axons was in the same range as previously measured average total occupancy of axons by mitochondria in mammals, e.g. ∼ 2% in cerebellar parallel fibers and optic nerve axons, ∼ 8% in olfactory receptor neurons, ∼ 6% in the fornix, ∼ 4% in retinal axons and; ∼ 3 to 9% in different axons of hippocampal axons (Perge et al. 2012; Perge et al. 2009; Faitg et al. 2021). Applying equation 8, longitudinal coverage = total occupancy/cross-sectional occupancy = 3.77/29.1= 0.130 or 13% on average.

### Mitochondrial increase in latency of action potential propagation between HVC and RA and its dependence on mitochondrial longitudinal coverage

We next investigated the impact of mitochondria on conduction velocity as action potentials propagate from HVC to RA. The longitudinal coverage of axons by mitochondria was varied in these simulations from 2 to 30%, in agreement with the 13% occupancy measured above as well as that of axons measured in other systems, and the known changes of mitochondrial coverage with age, injury and regeneration and activity (Faitg et al. 2021; Perge et al. 2009; Perge et al. 2012; Han, Baig, and Hammarlund 2016). The distance travelled by axons in a sagittal plane between HVC and RA was approximately 3 mm (Fig. 3B), a similar estimation to that done previously in another songbird, the zebra finch (Egger et al. 2020). Note that the distance travelled by axons may be larger when linking distal parts of HVC and RA that are not in the same sagittal plane.

The presence of mitochondria relative to ‘Gedankenexperiment’ mitochondria-free axons changed the amount of time required for an action potential to reach the RA for representative axonal sizes: for a typical 25% cross-sectional mitochondrial occupancy of a 0.4 μm non-myelinated diameter axon with 12.5% longitudinal coverage, action potentials accumulated an additional delay of 0.14 ms in their arrival to RA (Fig.3 E2) and a decrease in average conduction velocity of 2.1%. For smaller fibers with 0.2 μm in diameter, a typical 60% cross-sectional mitochondrial occupancy and 12.5% coverage, action potentials are delayed by 0.84 ms by mitochondria and have a decrease in average conduction velocity by 8.0%. Although the amount of additional delay depended on the axon diameter, the relative decrease in conduction velocity was similar for all diameters examined.

### Mitochondria induce anisotropic propagation of action potentials

The decrease in average conduction velocity along the pathway due to mitochondrial cross-sectional occupancy and longitudinal coverage is composed of small slow-downs as the action potential passes each individual mitochondrion. The amount of this local decrease depends on the properties of the mitochondrion. Hence the propagation velocity along an axon with constant diameter and biophysical properties is not expected to be constant but to vary as mitochondria are encountered.

Figure 4 shows an example of this anisotropy in action potential propagation in a simulated 0.3 μm diameter axon with 14% of the length containing 1 μm long mitochondria. The cross-sectional occupancy for each mitochondrion was drawn from a normal distribution with mean 0.3 and standard deviation 0.1. The figure shows the decrease in conduction velocity for each mitochondrion but doesn’t resolve the transition from mitochondrion free to mitochondrion containing axon sections.

**Figure 4.**
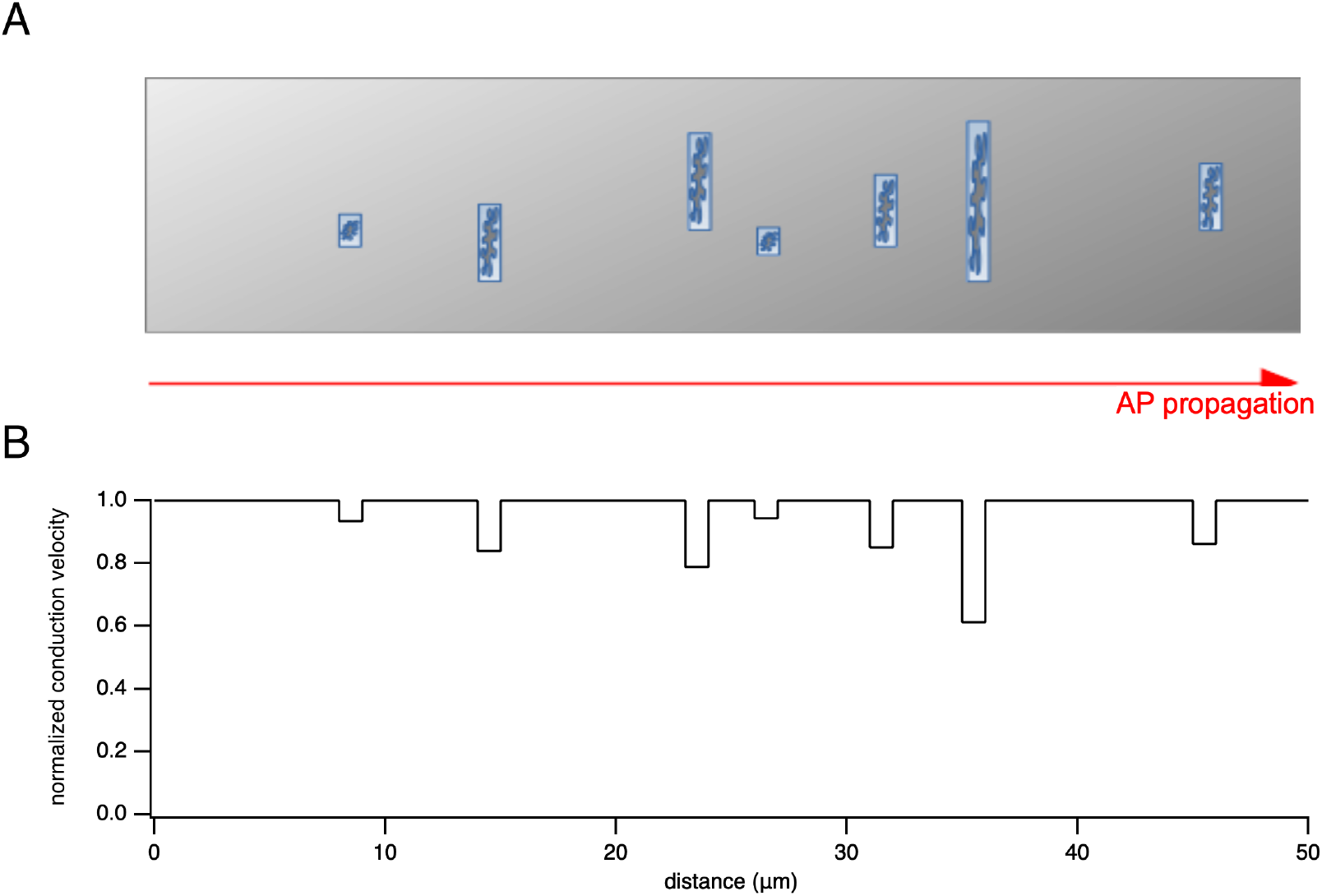
Model of anisotropic propagation of action potentials along axons. Schematic diagram illustrating the anisotropic propagation of action potentials along axons. **A**. Axon where the axon diameter has been disproportionately enlarged to accommodate the accurate cross section occupied by representiative mitochondria. **B**. Simulation. Due to the presence of mitochondria occupying part of the axonal volume, action potential propagation is locally delayed. Mitochondria have been drawn in panel A from a 0.3 μm diameter axon with 14% length containing 1 μm long mitochondria. The cross-sectional occupancy p was drawn from a normal distribution with mean 0.3 and standard deviation 0.1.

## Discussion

We present measurements of the size and density of mitochondria in unmyelinated axons from electron micrographs of axon bundles of the premotor pathway linking song nuclei HVC and RA from canaries. We found that mitochondria-containing axons range from ∼200 to 700 nm diameter. The canary motor pathway contains unmyelinated axons of small diameters that co-exist with myelinated axons in the bundles of axons running between song nuclei HVC and RA, as previously shown in the zebra finch (Egger et al. 2020). We found that the cross-sectional occupancy of axons by mitochondria depends on the axon diameter: occupancy is larger for smaller diameter axons.

Due to this reduction in the axoplasm by mitochondrial occupancy, our modeling showed that action potentials propagate anisotropically in axons, resulting in a local slow down of action potential propagation to about half its original speed for largest mitochondria-to-axon area ratios typically found in small axons. Whereas anisotropy of action potential conduction has been reported in myelinated fibers as ‘saltatory’ conduction, whereby conduction speed increases at the internodes, action potentials have been assumed to propagate homogeneously along a non-myelinated axonal branch of constant diameter. However we found that each time a propagating action potential meets a mitochondrion, the instantaneous conduction velocity can drop by 16% of its original value for an average 29% cross-sectional occupancy, and by up to 47% for the largest mitochondrial cross sectional occupancies (Fig. 2F). Interestingly, this phenomenon does not affect all axons alike, given the modest scaling of mitochondrial size with axon size. Indeed, mitochondrial cross-section occupancy shows an interesting phenomenon: small axons are more prone to a larger cross-section coverage, and thereby, to a larger local slow down of action potentials per mitochondrion. Why do mitochondria modestly scale with axon diameter? We can speculate that the size of mitochondria may be constrained by their evolutionary origin and their co-evolution within eucariotic cellular structures (Sagan 1967).

The non-uniform propagation of action potentials due to mitochondria may lead to additional nonlinearities when combined with features such as axonal tree branching and impedance mismatches (Manor, Koch, and Segev 1991), ion channel clustering, or leakage through axonal gap junctions (Alcami and El Hady 2019; Alcamí and Pereda 2019). Note that contrary to changes in axial resistance that also concomitantly change the surface of the plasma membrane, mitochondria locally change axial resistance without affecting plasma membrane capacitance or resistance.

The small axons found in the songbird premotor pathway linking HVC and RA are known to conduct a neural code that relies on submillisecond precision (Hahnloser, Kozhevnikov, and Fee 2002). Mitochondrial delays, that we estimate to be in the same time frame, thus likely constitute a design feature that the system has to take into consideration for its computations. Indeed, although submillisecond precision in the coordination of cell assemblies encoding song is necessary, a slight delay in conduction speed that delays the arrival of action potentials in RA within the millisecond range seems to pose a constraint that the system builds upon. Other features may constitute a stronger selective pressure to keep small unmyelinated axons in this pathway in the songbird brain. These may be related to miniaturization and volumetric constraints (Perge et al. 2009). However, the fast speed of conduction in other systems that strongly rely on submillisecond and fast conduction may be subject to a strong pressure whereby mitochondria-induced slowing down could not be tolerated since it would significantly alter the neural code necesssary for survival. An example of such system can be found in the auditory system, fine tuned for speed (Taschenberger and von Gersdorff 2000), which is formed by large myelinated axons from cells whose somata are found in the cochlear nucleus and whose axons project to the main nucleus of the trapezoid body (Grothe, Pecka, and McAlpine 2010). Since slowing down by mitochondria can be mostly overcome by larger axons, for which mitochondria scale sublinearly with their large size, and by myelination, which will favor a fast propagation of action potentials along neuronal and myelin membranes by their faster capacitive loading (Castelfranco and Hartline 2015; Alcami and El Hady 2019; Cohen et al. 2020), this combination of large and myelinated axons seems to be a solution that reduces action potential conduction delays.

Our results point towards an interesting paradox: on the one hand, energy, which as a first approximation can be estimated to be proportional to mitochondrial volume, is required to sustain axonal function, and in particular action potential generation and propagation (Borowsky and Collins 1989; Perge et al. 2009). Moreover, generating action potentials at higher frequencies leads to higher information encoding axons and likely requires larger mitochondrial volumes along axons (Perge et al. 2009). On the other hand, generating energy likely leads to a larger mitochondrial occupancy of the axoplasm, and thereby, to a slower conduction velocity. Thus, we postulate that there is a tradeoff between energy requirements for information coding by axons and conduction velocity. Interestingly, changes in metabolic demand or efficiency could lead to plasticity in mitochondria density and paradoxically to slowing down action potentials as a cost for higher energy supply.

Our computational model only explores resisitive changes in axial impedance, leaving aside contributions of mitochondrial capacitance. Our estimate of mitochondrial resistivity was consistent with the measurements of mitochondrial resistance of Jonas et al. (1999)(Jonas, Buchanan, and Kaczmarek 1999) and is likely to be an underestimate because of our assumptions about the dimensions of the resistor. A smaller resistor length would give a larger resistivity. However, increasing the mitochondrial resistivity would have only a minor effect on the results since the combined resistivity of the axoplasm and the mitochondrion saturates for larger values. The charge of mitochondrial capacitance seems to play a negligible contribution to the axial impedance change along the axon, since (Padmaraj et al. 2014) found little change in measured mitochondrial impedance for signals up to 10kHz, which is beyond neuronal computational time-scales, justifying our assumption. Finally, we assume a simple, cylindrical shape for a mitochondrion, which agrees with the low complexity index of mitochondria in hippocampal axons reported in (Faitg et al. 2021).

How widespread is the phenomenon reported here in axons across phylogeny? First, long small diameter axons are widely found in mammals (Perge et al. 2009; Braitenberg 1991; Wang et al. 2008) including humans (Aboitiz et al. 1992). Second, axonal occupancy by mitochondria seems to be in the same range as reported here in both invertebrates and vertebrates (Phelps et al. 2021; Perge et al. 2009), suggesting that the phenomenon is widespread across the phylogeny. If we consider the ∼ 0.1-0.8 ms delays in action potential propagation expected for 3mm long axons, and how these delays may generalize to axons one to two orders of magnitude longer, we would expect cumulated latencies in the millisecond to tens of miliseconds range, e.g. in the long non-myelinated fibers found in the human brain (Aboitiz et al. 1992). Remarkably, mitochondria show large levels of plasticity with activity age or disease (Faitg et al. 2021; Han, Baig, and Hammarlund 2016). It would be interesting to consider changes in action potential conduction velocity induced by these previously-reported changes in mitochondrial coverage.

The presence of additional organelles may analogously further contribute to delaying action potential conduction and increase the impact of organelles, here solely reduced to mitochondria. Finally, it would be interesting to model the impact of mitochondria in dendrites, which show a larger total mitochondrial occupancy than axons, e.g. in neocortical pyramidal cells (Lewis et al. 2018). There, mitochondrial occupancy may also interact with the propagation of synaptic events and dendritic spikes (London and Häusser 2005).

So far, cable theory has focused on biological cables free of organelles (Meunier and Segev 2001). Including intracellular organelles will add to our understanding of electrical signal propagation along biological cables found in neurons as well as in other cell types. Introducing mitochondria and more generally intracellular organelles as a structural design would allow to accurately model the propagation of action potentials, and more generally, of electrical signals.

## Acknowledgments

We thank Manfred Gahr for support and feedback throughout the study. We are thankful to Marianne Braun for the processing of samples for electron microscopy and advice throughout the course of this study, and animal caretakers at the Max Planck Institute for Ornithology for excellent care of canaries. We thank Wiebke Möbius and Moritz Hertel for advice on perfusions.

